# Single amino acid mutations effect Zika virus replication *in vitro* and virulence *in vivo*

**DOI:** 10.1101/2020.08.06.239392

**Authors:** Nicole M. Collette, Victoria H.I. Lao, Dina R. Weilhammer, Barbara Zingg, Shoshana D. Cohen, Mona Hwang, Lark L. Coffey, Sarah L. Grady, Adam T. Zemla, Monica K. Borucki

**Affiliations:** Biosciences and Biotechnology Division, Physical and Life Sciences Directorate, Lawrence Livermore National Laboratory, Livermore, CA, 94550, USA; Los Positas Community College, Livermore, CA, 94551, USA; Department of Pathology, Microbiology and Immunology, School of Veterinary Medicine, University of California, Davis, California, USA; Johns Hopkins University Applied Physics Laboratory, Maryland, USA; Computations Directorate, Lawrence Livermore National Laboratory, Livermore, CA, 94550, USA

## Abstract

The 2014-2016 Zika virus (ZIKV) epidemic in the Americas resulted in large deposits of next-generation sequencing data from clinical samples. This resource was mined to identify emerging mutations and trends in mutations as the outbreak progressed over time. Information on transmission dynamics, prevalence and persistence of intra-host mutants, and the position of a mutation on a protein were then used to prioritize 544 reported mutations based on their ability to impact ZIKV phenotype. Using this criteria, six mutants (representing naturally occurring mutations) were generated as synthetic infectious clones using a 2015 Puerto Rican epidemic strain PRVABC59 as the parental backbone. The phenotypes of these naturally occurring variants were examined using both cell culture and murine model systems. Mutants had distinct phenotypes, including changes in replication rate, embryo death, and decreased head size. In particular, a NS2B mutant previously detected during *in vivo* studies in rhesus macaques was found to cause lethal infections in adult mice, abortions in pregnant females, and increased viral genome copies in both brain tissue and blood of female mice. Additionally, mutants with changes in the region of NS3 that interfaces with NS5 during replication displayed reduced replication in the blood of adult mice. This analytical pathway, integrating both bioinformatic and wet lab experiments, provides a foundation for understanding how naturally occurring single mutations affect disease outcome and can be used to predict the of severity of future ZIKV outbreaks.

**Author summary:** To determine if naturally occurring individual mutations in the Zika virus epidemic genotype effect viral virulence or replication rate *in vitro* or *in vivo*, we generated an infectious clone representing the epidemic genotype of stain Puerto Rico, 2015. Using this clone, six mutants were created by changing nucleotides in the genome to cause one to two amino acid substitutions in the encoded proteins. The six mutants we generated represent mutations that differentiated the early epidemic genotype from genotypes that were either ancestral or that occurred later in the epidemic. We assayed each mutant for changes in growth rate, and for virulence in adult mice and pregnant mice. Three of the mutants caused catastrophic embryo effects including increased embryonic death or significant decrease in head diameter. Three other mutants that had mutations in a genome region associated with replication resulted in changes in *in vitro* and *in vivo* replication rates. These results illustrate the potential impact of individual mutations in viral phenotype.

## Introduction

Zika virus (ZIKV) is a mosquito-borne virus that replicates in both primates and *Aedes* mosquitoes (1). ZIKV belongs to the family *Flaviviridae* and genus *Flavivirus*, which includes other mosquito-borne and tick-borne human pathogens such as West Nile, dengue, yellow fever, and tick-borne encephalitis viruses. ZIKV was first discovered in 1947 in Uganda in a sentinel monkey and has circulated for decades in Africa and Asia. Phylogenetic studies have indicated that ZIKV has evolved into 2 major lineages: African and Asian/American, with reported differences in phenotype (2). Strains deriving from the Asian lineage emerged in Oceania and the Americas in 2013-2015, where they were associated with large outbreaks and rare cases of severe disease (3,4). Of particular concern was the appearance of devastating congenital abnormalities in babies born from infected mothers, characterized primarily by microcephaly. Hundreds of genomes were deposited during this outbreak, but analysis of ZIKV genomes from microcephaly cases revealed no conserved amino acid changes, suggesting that fetal abnormalities were not caused by an individual viral genetic feature (2,3,5,6). This finding resulted in increased research into the underlying mechanism of the microcephaly phenotype. We aimed to select, produce, and analyze the effects of, several mutations that appeared during the ZIKV outbreak.

ZIKV has a 10.7 kb genome of single-stranded, positive sense RNA that is translated into a polyprotein and cleaved into three structural proteins (capsid (C), precursor of membrane (prM/M), envelope (E)), and seven nonstructural proteins (NS1, NS2A, NS2B, NS3, NS4A, NS4B, and NS5). The structural proteins contribute to the formation of the mature infectious virion. Protein C forms the viral capsid after proteolytic cleavage inside the host cell, while prM/M and protein E are located on the viral envelope. prM is cleaved by host furin proteins to generate the mature M (membrane) protein. Protein E is the main constituent of the viral surface and is involved cell membrane fusion and endocytosis (7). The ZIKV non-structural proteins serve to regulate viral replication and host immune response. NS1 is a multifunctional glycoprotein that dimerizes in the membrane of the endoplasmic reticulum (ER) to participate in viral replication during late stages of infection. NS1 is also secreted in a hexameric form that contributes to immune evasion and pathogenesis (8,9). NS3 is important in viral assembly as a serino-protease that cleaves the viral polyprotein and requires NS2B as a cofactor. NS3 also participates in the viral replication complex as a helicase, a hydrolase and an RNA-triphosphatase. NS5 consists of methyltransferase and RNA-dependent RNA polymerase domains (10). Less is known about the functions of NS2A, NS4A, and NS4B, but all three are transmembrane proteins that form part of the viral replication complex on the ER membrane. All seven NS proteins have also been shown to interact with host proteins to dampen the interferon immune response and promote pathogenicity through diverse strategies. NS1 and NS4B, for example, inhibit interferon-beta production (11,12), while NS2B impairs the interferon-signaling pathway (12,13). NS5 has been found to bind and degrade human STAT2, which consequently blocks the interferon response (14,15).

To determine how mutations alter the infectious life cycle of ZIKV, both *in vitro* and *in vivo* model systems have been established. While wild-type mice have a robust Type I interferon response that prevents virulent infection by ZIKV due to the failure of NS5 to interact with the murine Stat2 protein (16,17), mutations in interferon receptors disable the interferon anti-viral response downstream of Stat2, leading to a virulent phenotype similar to what is seen in humans (15). One such mouse strain, *Ifnar1*^*-/-*^, has a knockout in the interferon α/β receptor (18,19). These animals are susceptible to infection and show recovery at clinically relevant doses of ZIKV (20). A second mouse model which is deficient in Type II interferon response, *Ifnα/β/γ/*^*-/-*^ *(AG129)* (18)(21), results in a much more severe infection with ZIKV (22). With both the α/β and γ interferon receptors mutated, this model recapitulates a lethal infection without any necessary changes made to the virus. Here, we utilize the *Ifnar* ^*-/-*^ (*a*.*k*.*a. Ifnar1* knockout, *Ifn α/β* ^*-/-*^, *A129)* mouse model to allow for survival and recovery using the wild-type virus in order to examine whether a given mutation (detected in the epidemic genotype) increases infection morbidity and/or mortality.

## Results

### Selection and synthesis of infectious clone genotype and mutants for analysis

Guided by computational analysis of ZIKV genotypes present in outbreak samples (23), six infectious clones of prevalent and persistent genotypes were synthesized and characterized for virulence using an A129 mouse model (17). Non-synonymous mutations that occurred at the interface of protein-protein interactions and those that were associated with microcephaly or highly virulent infections (10,23) were prioritized for reverse genetics studies. Mutations were also chosen based on their location within different proteins towards further defining their individual functions during viral replication and disease progression. Down-selected loci included polyprotein amino acid 123 (prM protein), polyprotein amino acid 894 (NS1), polyprotein amino acid 1404 (NS2B), and polyprotein amino acids 2074 and 2086 (NS3). A combined mutant genotype that had both the 2074 and the 2086 mutations (double mutant 2074/2086) was also selected to determine if changing both amino acids had an additive or synergistic effect on phenotype.

Two mutations, prM A123Vand NS3 H2086Y are reversions from the epidemic genotype to the ancestral amino acid (Malaysia 1966, strain P6-740, Accession # KX377336.1), while two others, NS1 G894A and NS3 M2074L, represent mutations that occurred during the course of the epidemic, as part of a relatively large clade (3,24). One mutation, NS2B, represents only 7 consensus sequences (as of July 2017) within the epidemic but is a high frequency variant that arises during *in vivo* infections of macaques and of mice dosed with interferon antibodies (23). In each case the detection of the mutation was associated with a change in the dynamics of the epidemic or a shift in the variant viral population within a host. The two NS3 mutants occur in the helicase domain of the protein, and in a region of NS3 predicted to interact with NS5 during replication (25).

### Generation of Synthetic Infectious Clone and Site-Directed Mutagenesis

The genomic sequence of the Puerto Rico 2015 strain (PRVABC59, accession # KU501215) was selected as the “parental” or wild-type (WT) genome as its sequence was obtained directly from a clinical specimen prior to passage in the laboratory (26), and because of its wide use in other ZIKV studies. The wild-type infectious clone was generated by inserting the full-length PRVABC59 cDNA sequence into vector pACNR1811 (27) based on the work of Tsetsarkin, et al. (28) and similar to Weger-Luccari et al. (29). The viral sequence was modified with intron sequences in NS1 and NS5, which reduce toxicity in *E. coli* cells following transformation (28). \ Site-directed-mutagenesis (SDM) was used to create the six independent infectious clones containing the mutation(s) found in Figure 1, and resulting clones were sequence verified. Additional mutations were attempted at other sites (NS1 Met 1404 Val, NS1 1143, NS5 2634 and 3328) but no mutants were produced at those sites despite repeated attempts, likely due to toxicity in *E. coli* or Vero cells.

**Figure 1.**
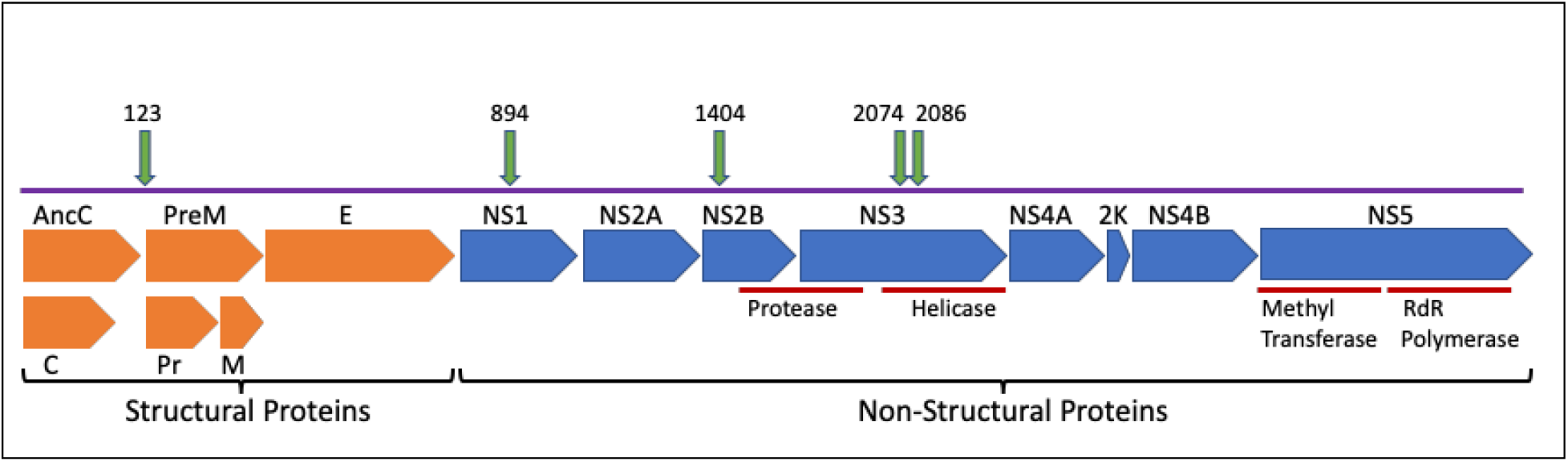
Schematic representation of the ZIKV genome. This representation shows viral proteins and locations of amino acid mutations chosen for in-depth examination in this study. Numbers indicate the polyprotein amino acid residue that was manipulated to generate synthetic mutants. orange= structural proteins; blue= non-structural proteins; green= mutation locations.

### *In vitro* assays

We assessed the phenotypic impact of individual mutations on the growth of the virus *in vitro* using a flow cytometry-based assay. Viral growth kinetics over time within Vero (primate) cells and C6/36 (mosquito) cells was first assessed using the WT virus. Viral growth, as assessed by the percentage of infected cells, peaked at 48 h in Vero cells and at 90 h in C6/36 cells (Figure 2A and B, respectively). We then assessed growth of mutant viruses at the peak time point in each cell line. In order to facilitate comparisons between individual experiments, data are represented as the fold change in the percentage of cells infected with mutant virus versus the percentage of cells infected with WT virus (Figure 2C and D). The mutants displayed varying growth characteristics in each cell line. Mutants 123 and 894 grew similarly as WT virus in Vero cells, but grew significantly faster in C6/36 insect cells. Infection with mutants 1404, 2074, and 2086 resulted in fewer infected cells at 48 hours in Vero cells. 1404 was the only mutant to result in a reduced number of infected C6/36 cells at 90 hours. Mutant 2086 was the only mutant in C6/36 cells that showed a similar rate of infection as WT at 90 hours; with the exception of mutant 1404, all other mutants showed higher percentages of infected cells at 90 hours post-infection. The 2074/2086 double mutant displayed significantly more growth versus WT virus in both cell lines. In order to validate the flow-cytometry based assay for the assessment of viral growth, we also assessed viral titer in the supernatant collected from infected cells at 48 and 90 hour time points, and found that plaque assay data closely correlates with flow-cytometry results. (Supplementary Fig. 1).

**Figure 2.**
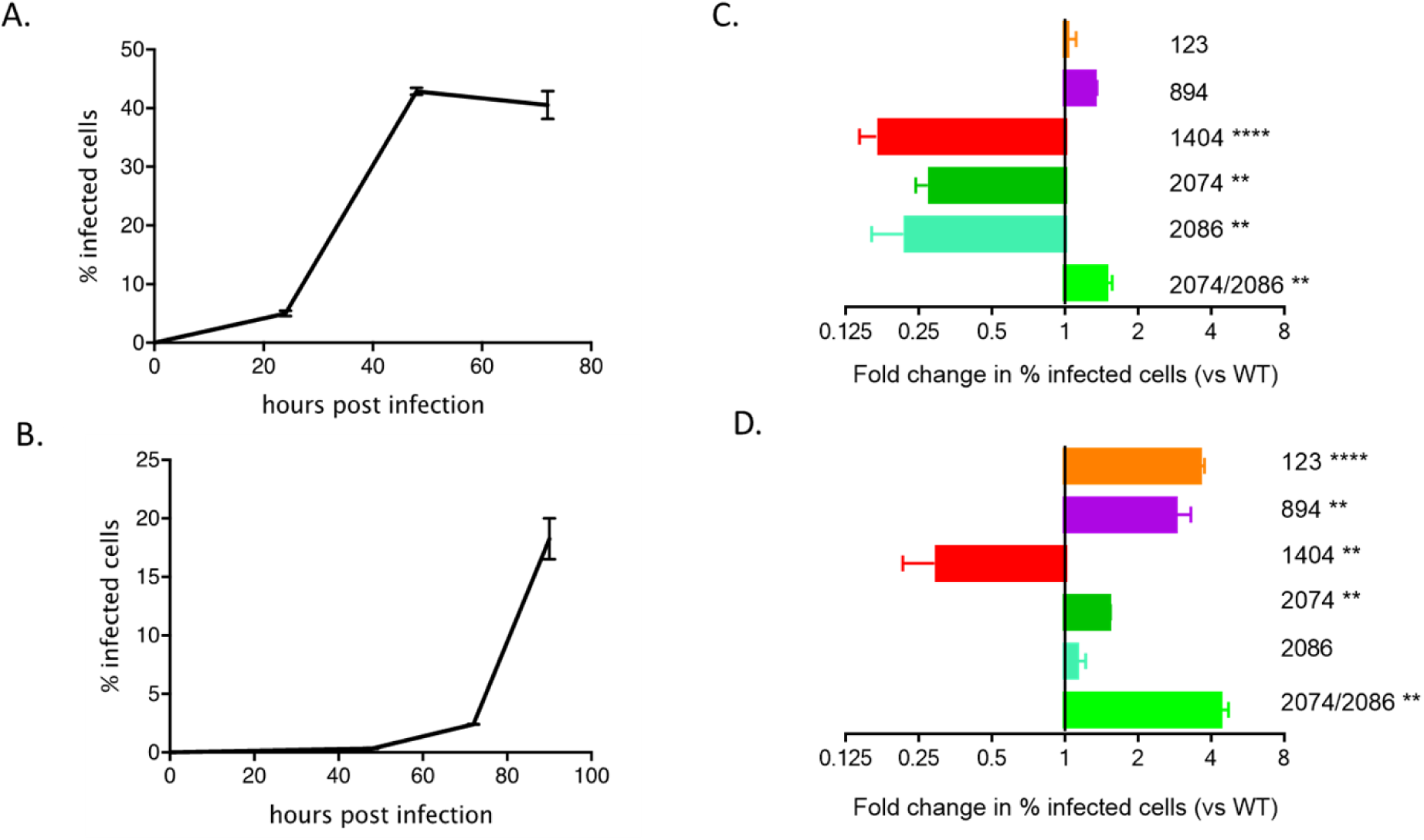
Mutants display varying growth characteristics in Vero and C6/36 cell lines. Vero and C6/36 cell lines were infected with WT or indicated mutant viruses at a MOI of 0.01 (Vero) or 0.05 (C6/36) and harvested for FACS analysis at the indicated time points. Representative time course plots are shown for the WT virus in Vero cells (A) and C6/36 cells (B). The fold change in infected cells observed with each mutant relative to WT are shown at the 48 h time point for Vero cells (C) and at the 90 h time point for C6/36 cells (D). **p<0.01, ****p<0.0001

### *In vivo* model

To determine if the WT infectious clone caused observable disease in the A129 mouse, groups of adult male and female *Ifnar1*^*-/-*^ mice were dosed with log 5 pfu of each virus subcutaneously in the hock location (30,31) and monitored for weight loss and neurological symptoms up to 14-21 days post-infection (dpi). As expected, WT infection led to weight loss (Figure 3), neurological symptoms (tremors, ataxia, and hindlimb paralysis), and persistence of viral genomes in tissues (brain, spleen, reproductive organs), consistent with other reports (17,20,31). When the infection was allowed to progress for 21 days, males had a significant reduction in testis size, as reported previously (20).

**Figure 3.**
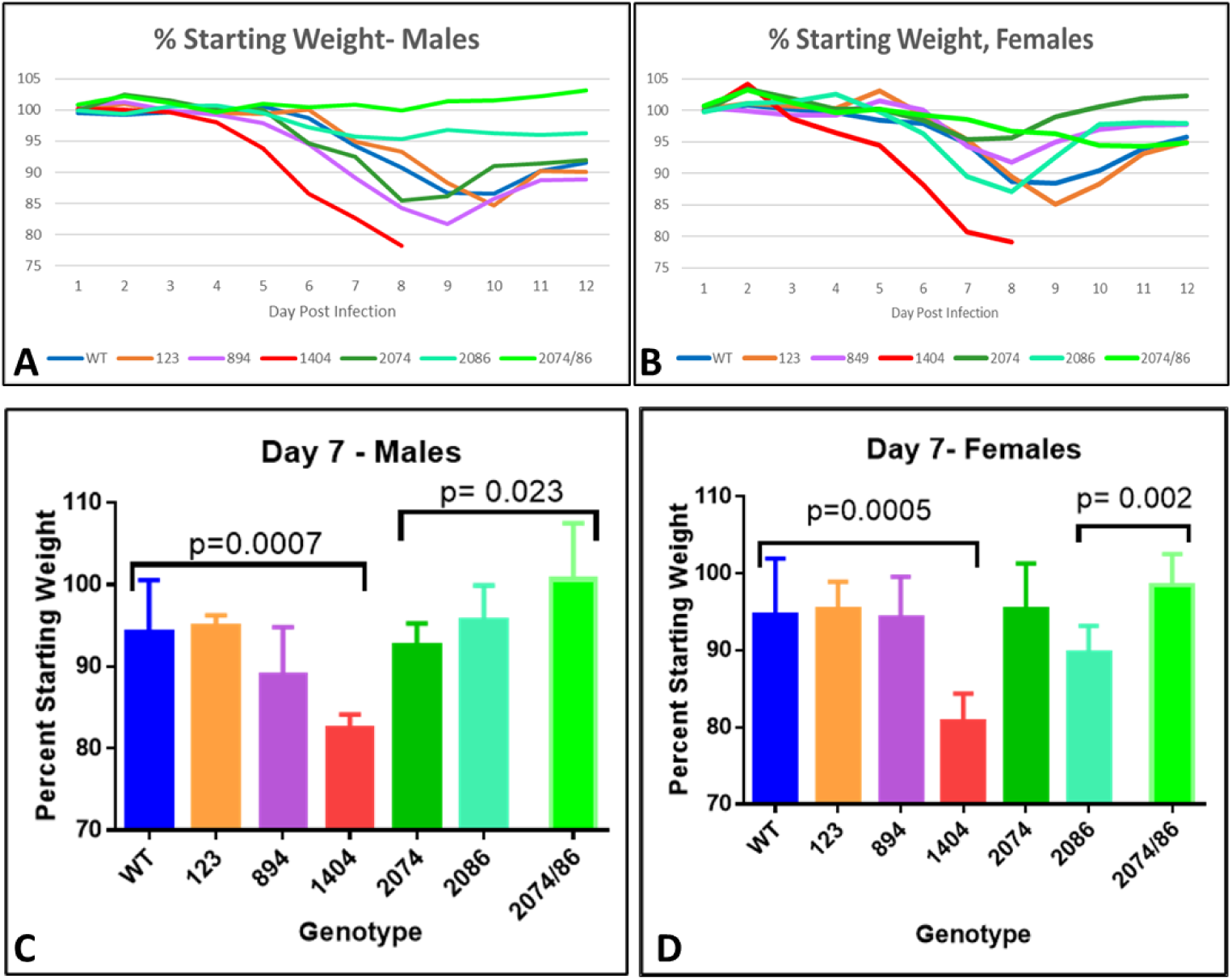
Weight loss and recovery after infection with synthetic ZIKV strains in adult mice. (A, B) Daily weight measurements in males (A) and females (B) measured as a percentage of starting weight. Mice of both sexes generally begin to regain weigh by 10 days post-infection. (C, D) Significant weight loss measured at day 7, compared to WT in males (C) and females (D). Values significantly different from WT are indicated by a bracket. P-values are indicated (Student’s T-test). Genotype 2074/86= 2074/2086

Infectious disease course and outcome for each mutant were next compared to that of the WT clone to determine if mice could recover from the infection and if viral RNA persisted in the tissues after recovery. Most mice developed symptoms at approximately 5-7 days post-injection. Symptoms included weight loss, mild dehydration, and progressed to neurological symptoms in some animals by approximately day 8-10, when symptom severity peaked for most mutants tested. Neurological sequelae that occurred late in infection included abnormal gait, disorientation, ataxia, and some animals also experienced temporary and reversible limb paralysis. A small number of animals experienced profound neurological symptoms such as seizures or loss of righting reflex and were euthanized.

Some differences in symptom severity, time of onset, and persistence were noted in some of the mutant strains. Males dosed with mutants 123 or 2086, both mutations that represented reversions to the Malaysia pre-epidemic amino acid sequence, became only mildly ill with the exception of one animal from each mutant group that required euthanasia. No animals in the 2074/2086 double mutant groups (male or female) needed to be euthanized, but hindlimb paralysis was observed at a frequency of one animal per group (n=6 per group). For mutant 894, euthanized animals exhibited dark, atrophied livers, and evidence of hemorrhagic symptoms with blood found in the intestines at necropsy. The most notable symptoms were observed in mice infected with mutant 1404; all became moribund within 7-8 days of infection and required euthanasia due to weight loss. Neurological symptoms were observed close to the euthanasia timepoint, consistent with the neurological symptom onset for other strains.

### Differences in mutant phenotype: Weight loss

Weight loss is often a robust clinical measure of disease progression in mice. Percent starting weight of mice infected with each mutant was compared to that of mice infected with WT to assess the overall impact of the infection on health (Figure 3). For mice infected with the WT infectious clone, peak weight loss occurred between day 9-10 in males, and day 8-9 in females. (Fig. 3A, B) with an overall maximum loss of 12-13%. Mice infected with mutant 123 lost approximately 15% by day 10 in males (Fig, 3A) and day 9 in females. Male mice infected with mutant 894 had a maximum weight loss of approximately 18% by day 9, while females experienced milder symptoms, with a peak weight loss of approximately 8% by day 8 post-infection, which may indicate a difference in severity between males and females.

Mice infected with mutant 1404 experienced the most dramatic weight loss, which was also observed more rapidly after infection as compared to WT. For this mutant strain, significant weight loss was observed starting at day 5 in males, (p= 0.0001, Fig. 3A), and slightly earlier, at day 4 in females (p= 0.047, Fig. 3B). By day 7 or 8 post-infection, all mice infected with 1404 had exhibited a large enough weight loss to warrant euthanasia (Fig. 3A, B), and was significantly more severe compared to WT in both males and females (Fig. 3C, D).

Mild infection was evident in female mice infected with mutant 2074 (Fig. 3D). These mice lost little weight, rebounded to above starting weight by day 14, and had less percent weight loss when compared to WT on day 9 (p = 0.051). However, male mice lost more weigh, close to 15% by day 8, potentially indicating gender-specific disease severity. Mice infected with mutant 2086, on the other hand, showed milder weight loss in males than females, with about 5% total weight loss in males, significantly less than WT at day 9 (p=0.023), and 13% loss in females. Male mice infected with mutants 2086 and 2074/2086 lost very little weight, less than 5% loss (Fig. 3A, C). However, females infected with 2074/2086 still had not regained weight by Day 14 (Fig 3B). Infection with combined mutant 2074/2086 exhibited the milder weight loss compared to either single mutant (p=0.023 in males, p=0.002 in females).

Overall, there was a trend of male mice showing increased weight loss and slower recovery to their starting weight as compared to females following infection. By day 14, when tissues were collected for analysis, all females except those infected with clone 1404 were within 3% of their starting weight, while males infected with any clone other than the WT weighed < 90% of their starting weight at the same time point. To determine whether this apparent sex-specific disease severity had anything to do with viral genomes present in male-specific tissue, the percent starting weight at day 14 was compared to the ZIKV genome copy number in the testes of each mouse. No correlation was found, suggesting the difference between male and female mice infected with the same virus could not be assigned solely to the testes acting as a reservoir for newly produced virus.

### Difference in pregnancy outcome

Pregnant female mice were used as a model to determine the effect of viral mutations on pregnancy and the development of microcephaly. Female mice were dosed in the hock after a successful observation of copulatory plug (embryonic day 4.5 (E4.5), Theiler Stage 6, (32)), consistent with the model used by Yockey, *et al*. (33). This mimicked infection during the first trimester of pregnancy in humans. Embryos were collected from presumptive pregnant females at E14.5 of pregnancy (Theiler Stage 23), which was also 10 days post-infection. For a minimum of 6 viable pregnancies per group, embryos were counted, and head diameters and crown-rump length were measured for each embryo with a pair of calipers after removal of membranes. Care was taken to collect individual embryo-placenta pairs for ZIKV genome copy analysis. Embryos and matching placentas were examined by RT-qPCR for viral genome copy number per μg tissue.

While there was no significantly different rate of pregnancy between any treatment group, a common phenotype for virus-infected pregnant females was loss of viable embryos or evidence of reabsorption. No loss of viable embryos was observed with PBS-dosed uninfected pregnant females, so our observations are unlikely to be a direct result of the injection procedure. Females that did not have any evidence of dead or live embryos at necropsy were not analyzed for pregnancy outcomes.

Pregnant mice infected with mutant 1404 aborted their litters and thus were not included in embryo measurements and pregnancy outcomes. All females were euthanized before 10-days post-infection due to weight loss and morbidity (E14.5). All embryos were lost from these pregnancies and no viable embryos were identified at necropsy. Reabsorbed embryos were observed in 5 WT-infected pregnancies. Compared to the WT strain, pregnant females infected with mutant 2086 showed similar embryo survival. However, mutant strains 123, 894, and 2074 had significantly fewer embryos survive as compared to WT (Fig. 4A). Notably, of all pregnant females for mutant 123, only one female had viable embryos at dissection at E14.5. There remaining females were pregnant but had deceased embryo numbers. Taken together, it indicates worse pregnancy outcomes for early pregnancy infection, particularly for mutants 1404 and 123, and to a lesser extent for mutants 894 and 2074, compared to WT.

**Figure 4.**
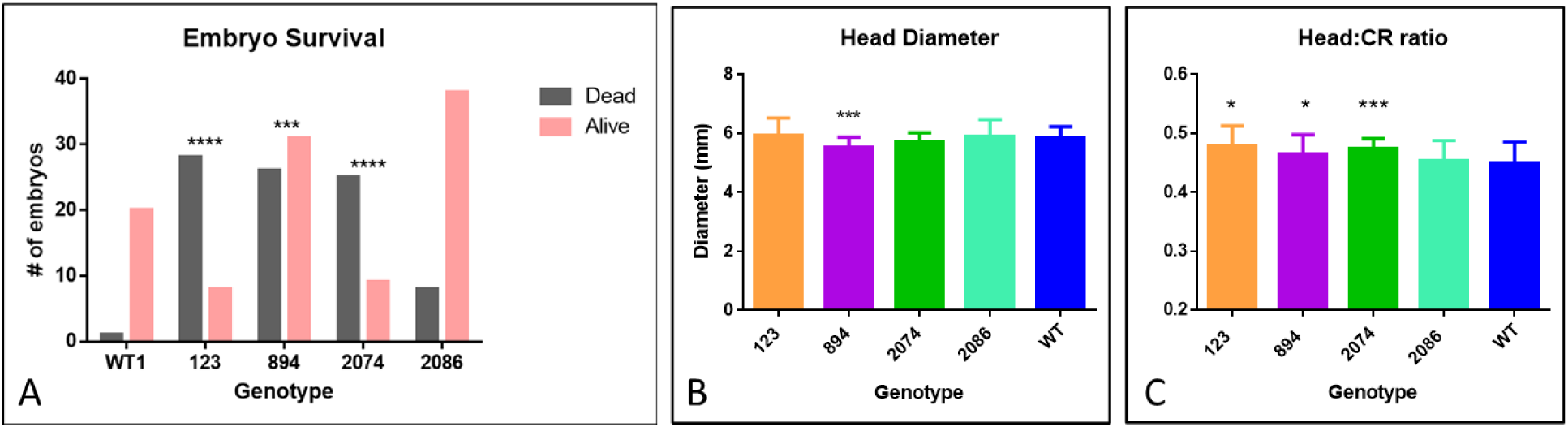
Embryo survival rates and embryo size measurements after infection with synthetic ZIKV strains. (A) total number of embryos recovered per pregnancy group. Gray-non-viable embryos, pink-viable embryos. A significant increase in number/percentage of non-viable embryos was found with mutants 123, 894, and 2074 compared to WT. (***= p<0.001, ****= p<0.0001). (B) Head diameter measurements compared to WT. Significantly smaller head diameter was noted in viable embryos of mutant 894 compared to WT (***=p<0.001). (C) Ratio of head diameter to crown-rump length; smaller ratios indicate smaller head diameters in relation to body size. (*= p<0.05; ***=p<0.001).

We observed a few embryos across the mutant strains that exhibited obvious microcephaly, defined as having notable brain loss, and we observed that these embryos also had affected eye development. More subtle differences in total embryo size or head diameter were observed in individual embryos. Microcephaly with eye deformities, and edema or failed closure of the neural tube were observed in pregnant females infected with the WT infectious clone (one embryo with microcephaly) and 4 embryos from pregnant females infected with mutant 2086 (3 with microcephaly and 1 with primary neural tube defects). Mutant strain 894 had a significant decrease in embryo head diameter (Fig. 4B), and the ratio of head diameter to crown-rump length of embryos was significantly decreased in pregnant females infected with mutant strains 123, 894, and 2074 (Fig. 4C). These metrics could be an indication of more severe growth retardation in the brains of the embryos from these pregnancies.

### ZIKV genome quantification in tissues by RT-qPCR

To examine tissue distribution, viral persistence post-infection, and viral copy number, brain, spleen, and testes or ovary tissues were analyzed for number of genome copies per μg total RNA using RT-qPCR. We analyzed tissues from males and non-pregnant females, in addition to pregnant females and embryos with placentas. Data from these analyses are shown in Figures 5, 6, and 7.

**Figure 5.**
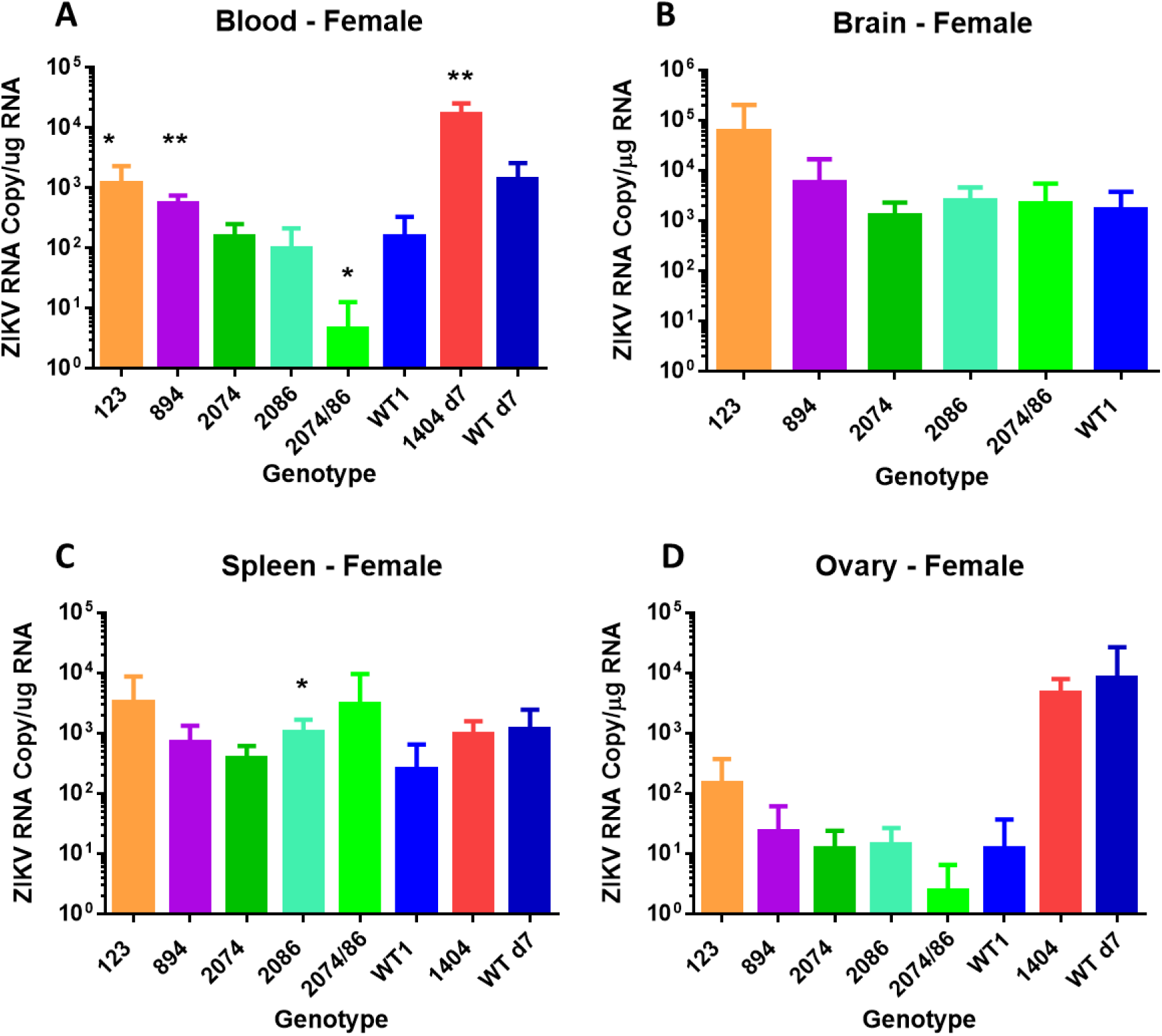
Viral copies of synthetic ZIKV strains in tissues of adult nonpregnant female mice at 14 days post-infection. Differences are compared to WT. Strain difference for mutant 1404 was compared to WT at day 7 since no animals remained at day 14 when the other samples were collected. (A) ZIKV genome copies identified per μg total RNA in whole blood. (* = p <0.05; ** =p <0.01). (B) ZIKV genome copies identified per μg total RNA from brain tissue. (C) ZIKV genome copies identified per μg total RNA in spleen. (* =p <0.05). (D) ZIKV genome copies identified per μg total RNA in ovary tissue. No significant differences were identified at day 14. Genotype 2074/86= 2074/2086

**Figure 6.**
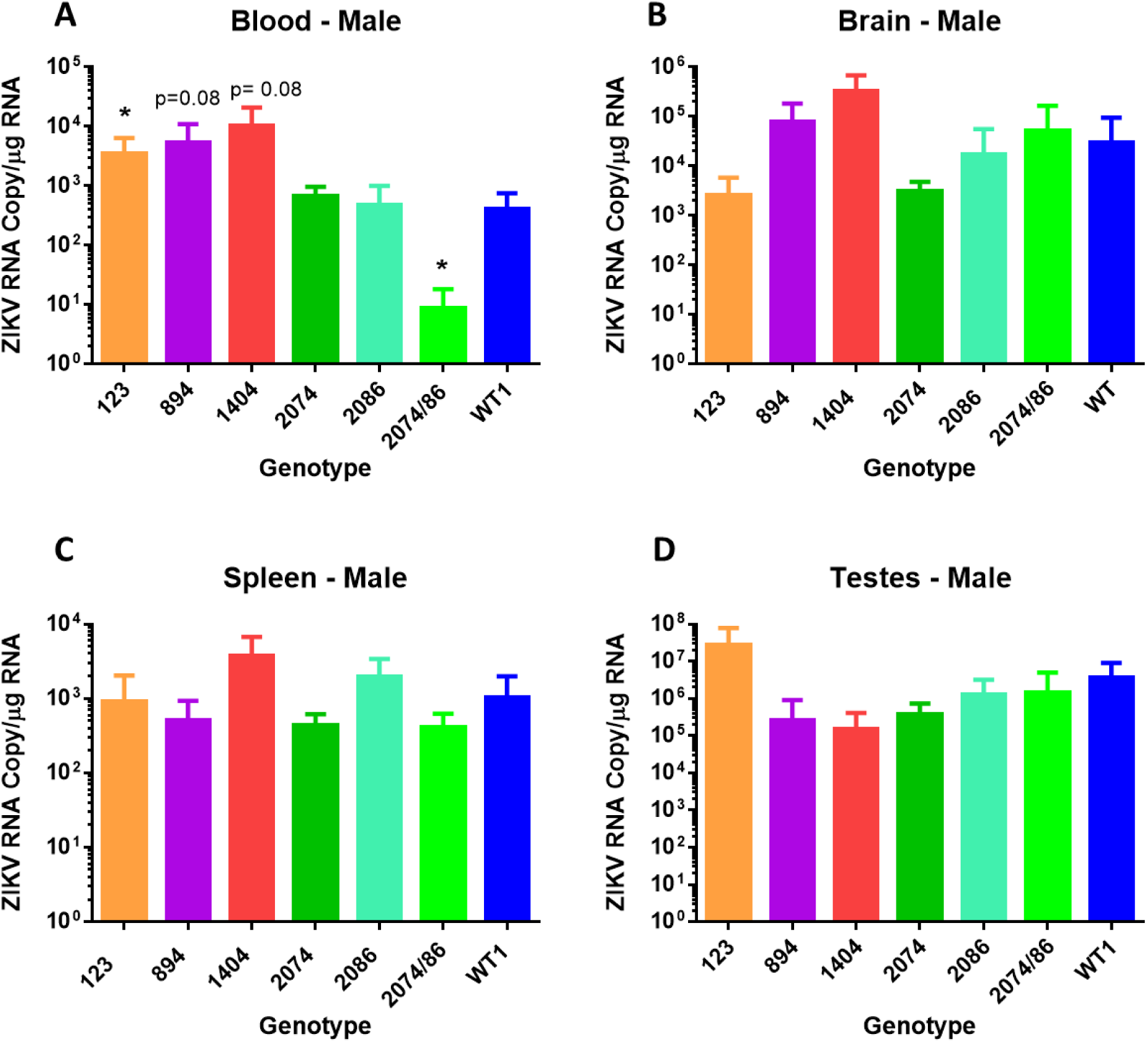
Viral copies of synthetic ZIKV strains in tissues of adult male mice. Differences are compared to WT. Strain difference for mutant 1404 was compared to WT at day 7 since no animals remained at day 14 when the other samples were collected. (A) ZIKV genome copies identified per μg total RNA in whole blood. (* = p < 0.05). (B) ZIKV genome copies identified per μg total RNA from brain tissue. (C) ZIKV genome copies identified per μg total RNA in spleen. (D) ZIKV genome copies identified per μg total RNA in ovary tissue. No significant differences were identified at day 14. Genotype 2074/86= 2074/2086

**Figure 7.**
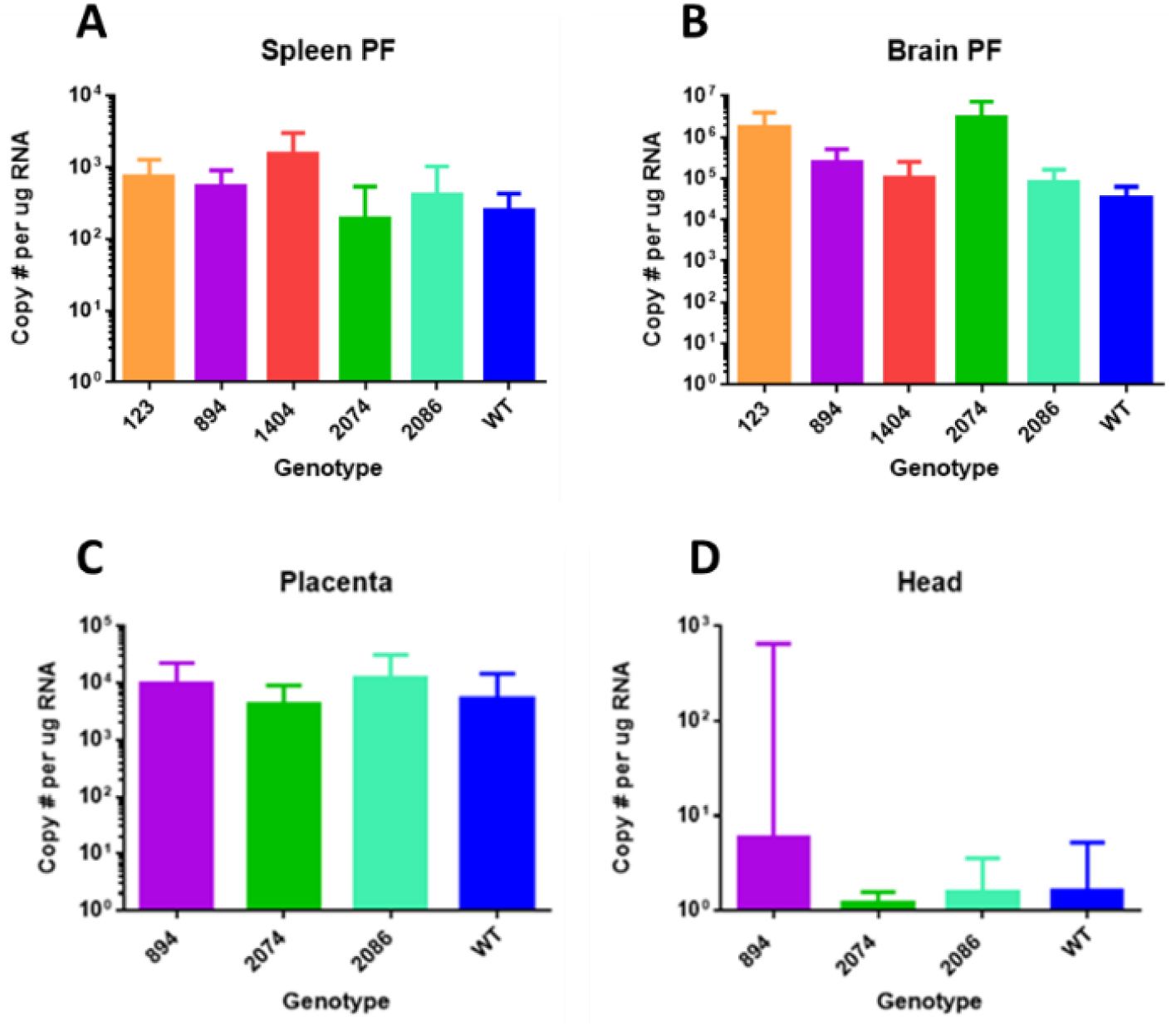
Tissues from pregnant mice (brain PF and spleen PF), embryos (head), and placentas were assayed via RT-qPCR for viral RNA. (A) ZIKV genome copies identified per μg total RNA in spleen of pregnant females. (B) ZIKV genome copies identified per μg total RNA in brain tissue of pregnant females. (C) ZIKV genome copies identified per μg total RNA in E14.5 placenta associated with embryos infected via mother at E4.5 of development. (D) ZIKV genome copies identified per μg total RNA in heads of E14.5 embryos infected via mother at E4.5 of development. No significant difference was detected between any of the mutants and the WT virus. In general, viral RNA was detected in the placenta but not brain tissue of embryos.

Female mice infected with mutants 123 and 894 had significantly higher blood viral titers as compared to mice infected with WT (Fig 5A), as did female mice infected with mutant 1404 when compared to females infected with WT at day 7. The blood titer of the double mutant 2074/2086, however, was significantly lower than in WT samples. In general, there was a trend of mutants 123 and 894 to have the highest ZIKV genome copies in infected female mice in all tissues (Fig. 5A-C). The major exception was the spleen, where mutant 2086 had the highest titer (Fig. 5C).

As mice infected with mutant 1404 required euthanasia by day 7 or 8 (Fig. 2A, B), tissues from this group were collected approximately 6-7 days earlier than other groups. To allow for direct comparison between 1404 mice and WT mice, an additional group of females were infected with WT virus and euthanized at day 7. Female mice infected with mutant 1404 had an increased ZIKV genome copy number in brain tissue as compared to female mice infected with the WT infectious clone at day 7 (Fig. 5B).

Similar to the female mice, male mice infected with mutant 123 had significantly elevated viral genome copies in the blood, and mice infected with the double mutant 2074/2086 had significantly lower blood levels of viral RNA (Fig 6A). There was no statically significant difference observed in any of the other tissues. It should be noted, however, that the ZIKV genome copy number of each mutant in the testes was extremely high, especially for mutant 123 which had a titer of log 7 in infected male mice, consistent with the findings of other groups (20,34). This is indicative of the ongoing risk for sexual transmission of ZIKV (35).

While some mutants behaved similarly in both sexes, others showed sex-specific differences in viral genome copy number. For example, mice infected with mutant strain 894 showed a higher level of viral RNA in the blood of males compared to females, which is consistent with the sex differences observed in weight loss (Fig. 2). Low copy numbers of ZIKV genomes in the blood from mutant 2074/2086 infected mice of both sexes is consistent with the very mild weight loss observed (Fig. 2). Very low viral RNA copy number identified in ovary tissues of infected female mice compared to testes of male infected mice may indicate that this tissue is not a significant source of possible recurring infection for females with subsequent pregnancies.

When embryos were collected at E14.5 for measurements and viral copy number, tissues from the pregnant females (day 10 of infection) were also collected. At this time point, we did not observe any statistically significant differences in viral copy number compared to WT ZIKV genome copy number in spleen or brain of pregnant females (Fig. 7A, B). These copy numbers were also not appreciably different from viral genome copy observed at day 14 in non-pregnant females (Fig. 5B, C). Analysis of ZIKV genome copy number in embryo-related tissues, placenta and embryonic heads, revealed much higher viral copy number in placental tissue compared to the embryo itself (Fig. 7C, D). The ZIKV genome copy number in embryos varied substantially from embryo to embryo, despite consistent viral genome copies in maternal tissues and placentas. This may reveal some stochastic mechanism for susceptibility of transmission to the embryo, in which some embryos are more severely affected than others, even within the same litter. Placenta tissues (Fig. 7C) revealed significant viral copy number present even after organogenesis is complete, suggesting a potential ongoing reservoir for infection throughout gestation.

### Testing of tissue samples for reversion to wild type

Due to the high mutation rate of RNA viruses such as ZIKV, mutations established using SDM may revert back to the WT amino acid as the virus replicates within the host, especially in cases where the mutation is less fit. For each mutant, with the exception of double mutant 2074/2086, 16 to 18 tissue samples that had been assayed by RT-qPCR were tested for by Sanger sequencing for reversions of the mutation to the WT amino acid (Suppl Table 1, Suppl Fig. 2). No reversions in any of the tissue samples were detected for mutants 123, 894, and 1404. One of 17 tissue samples from mutant 2074 (female, blood), and 1of 18 the 2086 samples showed total reversion (1 male spleen sample). Four more of the 2086 samples showed partial reversion as shown by multiple peaks at the mutation site (1female spleen, 3 placenta samples). This result is not unexpected given that both the 2074 and the 2086 mutations are located in a region of the NS3 protein predicted to play a role in replication. Both of these mutants grew significantly more slowly in Vero cells as compared to WT (Fig. 2). For mutant 2086, the mutant amino acid was that of the ancestral, pre-epidemic genotype so these data indicate that the WT epidemic amino acid, H2086, increased the replication rate of the epidemic genotype.

## Discussion

Single amino acid mutations have been shown to have significant impact on ZIKV infection phenotypes (11,36,37), as well as diseases caused by other mosquito-borne viruses such as West Nile and chikungunya (38,39). Many mutations were identified during the course of the ZIKV outbreak in the Americas (3,28,40), and we selected several to characterize based on their transmission dynamics, protein structure models, and deep sequencing variant data. In total, six infectious clone-derived mutants were created and analyzed for changes in virulence using *in vitro* and *in vivo* assays.

For the mutants that reverted to the ancestral Malaysian strain amino acid residue, prM A123V and NS3 H2086Y, we anticipated that virulence would be attenuated as compared to the WT epidemic clone. Similarly, we anticipated that changes that occurred during the outbreak and spread extensively, including NS1 G894A and NS3 M2074L, would be either neutral or beneficial to transmission (*i*.*e*., increase viral genome copy number in blood, increased replication rate, etc.). Finally, given that NS2B M1404I increases in frequency in the pregnant macaque model (23,41) we anticipated that the pregnancy and embryo physiological environment is conducive to replication of virus, and thus this mutation may be associated with poor pregnancy outcome.

The ancestral Malaysian strain has a Val at residue 123 whereas the WT epidemic strain is characterized by an Ala at this site (3,42). SDM was used to change the Ala in the WT infectious clone to a Val. Polyprotein residue 123 (PrM protein residue 1) is located near residue 139 (PrM residue 17) in the native folded protein, and a S139N mutation at this site has been shown to contribute to microcephaly (36). The first 40 residues of the PrM protein are a hot spot for mutations that distinguish the epidemic strain from pre-epidemic strains and may affect protein structure and interactions (43).

Given that ZIKV has only recently been associated with microcephaly, we anticipated that reverting residue 123 to the ancestral Malaysian amino acid might result in attenuation of the virus in the mouse model. Unexpectedly, the mutant infection resulted in increased embryo death as compared to WT (Fig. 4A). In fact, none of the embryos survived in 4 of the 6 litters. It is possible that this ancestral mutation causes fetal death more quickly during human pregnancy as well, potentially resulting in fetal loss that remains unnoticed if it occurs early in pregnancy. A loss of fetus early in pregnancy would be unlikely to be associated with microcephaly, as measurements of brain size would not be performed. Interestingly, mutant 123 infection also caused increased viral genome copies in the blood of male and female mice as compared to mice infected with the WT infectious clone (Fig. 5A, 6A), and grew significantly faster in mosquito cell line C6/36 (Fig. 2B, D). It seems reasonable, then, that this mutation, when in the context of the 2014-16 epidemic genotype, actually increased virulence and the potential for transmission.

The Gly to Ala mutation at residue 894 was first detected in 2015 in Guatemala, Honduras, and Mexico and spread extensively in 2016-17 to multiple additional countries. This mutation was selected for study due to its extensive transmission, persistence, and association with virulent disease, including multiple microcephaly cases and a fulminant infection that was notable due to its extraordinarily high viremia and its nonsexual transmission between individuals (44). Protein structural modeling places residue 894 on the surface of the NS1 protein, and while the WT glycine residue is neutral, alanine is hydrophobic. NS1 plays an essential role in viral replication and immune evasion (6), and residue 894 falls in the alpha/beta wing domain, a region of NS1 that may interact with the host immune response (45). Together, this information suggested that this mutation could have a significant effect on protein structure and function.

Mutant 894 infection resulted in increased viral genome copies in the blood of both male and female mice (Fig. 5A, 6A), and significantly increased embryo death (Fig. 4A). Mutant 894 also grew more quickly in Vero and C6/36 cells as compared to WT (Fig. 2). The NS1 protein is known to alter host interferon response (6,11,45), but in our system, both Vero host cells and *Ifnar1*^*-/-*^ mice are interferon-deficient. Interestingly, even when removing the variable of host interferon response, mutant 894 still appeared to positively effect viral yields. This suggests that the 894 mutation increases virulence in an interferon-independent manner. Further work would be necessary to determine if it also improved virulence by dampening host interferon response.

Mutant M1404I, located within protein NS2B, was selected for analysis because it was a high frequency variant in animal models (23), although it was not associated with epidemic spread. In the present study, mutant 1404 was highly virulent in both adult mice, pregnant mice, and embryos. Early, lethal weight loss was observed in both adult male and female mice, and abortion in pregnant females (Fig. 3B, 3D, 4A). Compared to female mice infected with the WT strain, animals infected with mutant 1404 had increased viral genome copies detected in brain tissue and in blood of female mice (Fig. 5A, B).

Interestingly, mutant 1404 grew more slowly than WT in both Vero cells and C6/36 cells (Fig. 2). If clinical samples containing this mutation are amplified in Vero or C6/36 cells prior to sequencing, the delayed growth in these cell lines may provide an explanation for the fact that 1404 Ile is seldom found ZIKV genome consensus sequences. It is possible that the lower yields of this virus from tissue culture cells could be in part due to some level of cytotoxicity. One intriguing recent study suggests that NS2B-NS3 protease complex induces cleavage of septins during the cytokinesis stage of mitosis, and aberrant NS2B-NS3 activity can proteolytically cleave septins, resulting in failed cytokinesis, and abundant cell death (46). Thus, it is possible that the NS2B M1404I mutation may enhance this off-target cleavage in cells and increase cell death during ZIKV infection. Further studies focusing on the mechanism of pathogenesis associated with this mutation are warranted.

The NS3 Met 2074 Leu mutation was present in ZIKV genotypes circulating in Guatemala, Honduras, Nicaragua, and Mexico and was associated with several microcephaly cases as well as a highly virulent infection (47), thus we hypothesized this mutation might increase the viral replication rate and increase pathogenicity. Mutant 2074 grew significantly more slowly in Vero cells but outgrew the WT infectious clone in C6/36 cells at all timepoints (Fig. 2). Thus, it is possible that this mutation increases growth rate in mosquitoes, although determination of growth rate in live mosquitoes would be necessary to verify this notion. Interestingly, although mutant 2074 grew more slowly in nonpregnant female mice (Fig 5), it caused greater embryo death as compared to WT (Fig. 4) indicating increased virulence in embryos despite decreased replication rate.

The NS3 H2086Y mutation was a reversion to the non-epidemic, ancestral amino acid. We thus hypothesized that it would result in an attenuated phenotype. *In vitro* and *in vivo* testing both supported this hypothesis. The mutant had a decreased replication rate in Vero cells (Fig. 2) and caused less severe disease in adult male mice as compared to WT (Fig. 3). Male mice infected with mutant 2086 and had significantly less weight loss and overall had milder symptoms of infection. The number of viral genome copies in the spleen of female mice was reduced significantly compared to WT (Fig. 5) and importantly, mutant 2086 was the only mutant not to increase embryo death as compared to WT (Fig. 4).

A NS3 M2074L and NS3 H2086Y double-mutant was generated to determine if the mutations had a combinatorial effect on *in vitro* or *in vivo* phenotypes. In total, we tested four combinations of amino acid residues at these two sites. The WT infectious clone has 2074 Met and 2086 His, which represents the most common epidemic genotype. The single-mutant M2074L represents a genotype that emerged later in the epidemic and which was widely transmitted. The single-mutant H2086Y reflects the ancestral strain. The double-mutant includes the late epidemic 2074 Leu and the ancestral 2086 Tyr. Both locations are present on the face of NS3 (23). Unexpectedly, the double-mutant grew more rapidly in both Vero and C6/36 cell lines as compared to the WT (Fig. 2) but appeared to replicate relatively poorly *in vivo* as the number of viral genome copies was decreased in blood of both male and female mice (Fig. 4, 5). Male mice lost little weight and no mice infected with the 2074/2086 mutant required euthanasia (Fig. 3), which is consistent with the lower viral copy number identified in blood. These data indicate that the different combinations of mutation at 2074 and 2086 do impact viral growth rate differently, although no clear pattern of synergy was observed.

## Conclusion

In this study, an infectious clone representing an epidemic-associated consensus sequence of ZIKV was generated and used as a template for site-directed mutagenesis. Several mutations were selected based upon sequence database information, deep sequencing data, and protein structure analysis (23), and the associated infectious clones were generated and assayed for changes in *in vitro* and *in vivo* model systems. Mutations that chosen for further study were distributed throughout the genome, with the exception of two mutations (2074 and 2086) that were closely linked in the NS3 genomic region.

Based on our *in vitro* data, we conclude that mutations do not always similarly effect replication rates in insect and mammalian cells. Mutations at locations 1404, 2074, and 2086 reduced replication fitness in mammalian cells, while only 1404 reduced replication fitness in mosquito cells. Mutations 123, 894, 2074, and 2074/2086 may improve fitness in the mosquito vector in the wild and contribute to the persistence or spread of these mutations during the epidemic and future outbreaks.

Importantly, our *in vivo* data showed that mutants 123 and 1404 had catastrophic embryo effects when infection was introduced during embryonic development, and additionally, mutant 1404 proved to be highly virulent and caused host death in adult mice as well. While mutant 894 was not as catastrophic for embryo survival, it did cause increased embryonic death and also a significant decrease in head diameter and head diameter:crown-rump length ratio, suggesting persistent effects during embryonic development and potential contribution to Congenital Zika Syndrome after birth.

NS3 mutants 2074 and 2086, along with the combined mutant 2074/2086 resulted in detrimental effects on replication in Vero cells, with improved replication in mosquito cells. These mutations both reside in the helicase domain of NS3 (25), suggesting that viral helicase activity might be differentially regulated in insect and mammalian cells.

In summary, we have identified differential effects *in vitro* and *in vivo* of specific point mutations in the ZIKV genome that may be predictive of the severity of future ZIKV outbreaks when their persistence in the environment can be identified.

## Methods

### Virus

ZIKV strain PRVABC59 (Human/2015/Puerto Rico) was obtained from BEI Resources (NR50240 ZIKV). The complete genomic sequence of PRVABC59 has been determined (GenBank Accession: KU501215). The PRVABC59 virus stock was passaged a total of 5 times in Vero cells prior to sequencing according to the GenBank description of KX087101. Stock virus was separated from cells and debris by filter concentration and/or ultracentrifugation (48). This viral stock was used as our initial control strain for *in vitro* and *in vivo* infections.

### Construction of infectious clone

The PRVABC59 infectious clone was generated based on the methods described in (28). The original infectious clone contained the sequence corresponding to the Paraiba_01/2015 strain of the virus inserted into a low-copy plasmid backbone, pACNR1811 (27). The Paraiba_01/2015 sequence was removed and the PRVABC59 sequence was synthesized and cloned into the linearized backbone using Gibson assembly. The PRVABC59 sequence contained two intron sequences in the NS1 and NS5 genes to nullify toxicity during propagation in *E. coli*, as described in (28). The resulting clone was transformed into MC1061/P3 competent *E. coli* and sequence verified. Virus stocks were generated by transfecting the clone into Vero cells using Lipofectamine 2000 (Thermo Fisher, cat# 11668-019), and harvested at various times post-transfection.

### Site directed mutagenesis

Using the PRVABC59 synthetic infectious clone as a template, point mutations were generated at nucleotides 123, 894, 1404, 2074, 2086, and the double mutant 2074/2086 of the genomic ZIKV sequence by SDM. Q5 Site-Directed mutagenesis kit (New England BioLabs) was used to generate point mutations in the WT infectious clone. Primers for each mutant were designed using NEBaseChanger(tm). Q5’s Quick protocol was modified to improve the transformation efficiency for WT plasmid (14kb). A gradient PCR with +/-5°C adjustment to predicted annealing temperature was performed for each mutant. PCR amplicon was purified with E-Gel EX gel (Invitrogen) and QIAquick PCR Purification Kit (Qiagen). A minimum of 50 ng of purified PCR product was used in the Kinase, Ligase and DpnI treatment reaction. Transformants were size verified by PCR using primers designed to flank the region containing mutation. The entire length of the WT mutant was sequenced by Elim Biopharm. Primers used in mutagenesis are shown in Suppl. Table 2.

### *In vitro* studies

Viral growth *in vitro* was assessed using confluent Vero or C6/36 monolayers in 96-well plates following inoculation with a multiplicity of infection (MOI) of 0.01 (Vero) or 0.05 (C6/36) of each mutant or wild type virus stock in triplicate. Cells and virus were incubated for 1 h, then inoculum was removed, and monolayers were washed three times with PBS and overlaid with growth media. Cells and supernatants were harvested at 24, 48, and 72 h (Vero) or 48, 72, and 90 h (C6/36) post-infection. Cells were processed for intracellular flow cytometry staining using BD Cytofix/Cytoperm (BD Biosciences) according to manufacturer’s instructions. Cells were incubated with mouse anti-Zika antibody 4G2 (Millipore) at 1:500 dilution for 60 min at room temperature, followed by goat anti-mouse AF488 (Molecular Probes) at 1:1000 dilution for 30 min at room temperature. Flow cytometry was performed using a FACS Aria Fusion and data were analyzed using FlowJo software. The number of plaque forming units (PFU) in the collected supernatants was assessed by standard plaque assay on Vero cells at the 48 h (Vero supernatants) and 90 h (C6/36 supernatants) time points.

### RNA extraction

For RNA extractions, whole blood or tissue were used to isolate total RNA from each sample. Whole blood that had been collected in potassium EDTA was mixed with Trizol LS (Thermo Fisher) and isolated according to manufacturer’s instructions. Tissues were isolated at the time of necropsy and stored in RNALater (Invitrogen) at −20°C. Tissue was removed from RNALater and homogenized using a BeadBug(tm) bead homogenizer with 3mm ceramic beads in 600ul RLT buffer with beta-mercaptoethanol added. Tissue RNA from male and nonpregnant female was isolated with Qiagen RNEasy columns (Qiagen), with on-column DNase I digestion (Qiagen). Tissue isolations from pregnant mice were performed using a QIAcube Connect, with the standard protocol for tissue isolation. Isolated RNA was stored at −80°C until RT-qPCR analysis.

### RT-qPCR

Viral RNA was reverse transcribed and amplified using the SuperScript(tm) III One-Step RT-PCR System (Thermo Fisher Scientific) following manufacturer’s instructions. RT-qPCR conditions consisted of 98°C for 30 s, followed by 35 cycles of 98°C for 10 s, 60°C for 20 s, and 72°C for 1 min. The final cycle was 72°C for 2 min. Primers ZIKV 835, ZIKV 911, and probe ZIKV 860 from Lanciotti et al. (49) were used to determine number of genome copies per ug RNA. RNA template was synthesized (Genscript) and used for the standard curve.

### Mouse infection

These studies were carried out in strict accordance with the recommendations in the Guide for the Care and Use of Laboratory Animals and the National Institute of Health. All efforts were made to minimize suffering of animals. All animals were housed in ABSL2 conditions in an AAALAC-accredited facility, and the protocol was approved by the LLNL Institutional Animal Care and Use Committee (IACUC), which includes ethics in evaluation of protocols. Inbred *Ifnar1*^*-/-*^ mice (IFNAR1 knockout, A129, B6(Cg)-Ifnar1^tm1.2Ees^/J, Jax stock number 028288) were bred in-house and maintained in positive-airflow barrier-housing under specific pathogen-free conditions. Animals were moved into negative airflow ABSL2 containment housing 24 hours prior to ZIKV infection. For males and non-pregnant females, ZIKV infection (PRVABC59 strain and synthetic mutant clones) was injected subcutaneously/intra-dermally into the hock region of the animal (30) at 1 × 10^5 plaque forming units (PFU). Mice were observed at least daily with body condition scores and animal weights measured. Animals that reached the humane endpoint were euthanized. At up to 14 days post-infection, tissues were collected immediately after euthanasia (brain, spleen, and testis or ovary) and stored in RNA-Later (Qiagen) according to manufacturer’s instructions until RNA isolation and RNA analysis. For pregnant females, *Ifnar1*^*-/-*^ mice were mated, and embryonic day 0.5 (E0.5) of gestation was considered to be noon on the day a copulatory plug was observed. Females were infected via hock injection at E4.5 of pregnancy. At E14.5, embryos were measured (head diameter and crown-rump length), and both embryo (head, body, placenta with attached membranes) and female tissues (brain, spleen, ovary and pregnant uterus) were collected and stored in RNAlater. Blood was also collected from adult animals and stored in Trizol LS (Invitrogen) until RNA extraction. RNA was extracted from mouse tissues using the RNeasy Mini Kit (Qiagen) according to manufacturer’s instructions, after being homogenized in lysis buffer using 1.5 and 3.0 mm Zirconium Lysis BeadBug(tm) beads and a BeadBug 3(tm) homogenizer (Benchmark Scientific).

## Acknowledgements

This work was performed under the auspices of the U.S. Department of Energy by Lawrence Livermore National Laboratory under Contract DE-AC52-07NA27344. This study was supported by the Office of Research Infrastructure Programs/OD (P51OD011107), start-up funds from the University of California, Davis School of Veterinary Medicine Pathology, Microbiology and Immunology Department to L.L.C

## Author contributions

Conceptualization: M.K.B., N.M.C., S.L.G. Methodology: M.K.B., N.M.C., S.L.G. Investigation: N.M.C., M.K.B. V.I.L., S.L.G., A.T.Z., M.H., .S.D.C., D.R.W. L.L.C. Writing-original draft: N.M.C, M.K.B., S.L.G. Writing-review and editing: N.M.C, M.K.B., S.L.G. D.R.W. Visualization: M.K.B., N.M.C., S.L.G., A.T.Z. Supervision and project administration: M.K.B. Funding acquisition: M.K.B.

